# Accurate cell type deconvolution in spatial transcriptomics using a batch effect-free strategy

**DOI:** 10.1101/2022.12.15.520612

**Authors:** Linhua Wang, Ling Wu, Chaozhong Liu, Wanli Wang, Xiang H.-F. Zhang, Zhandong Liu

## Abstract

Sequencing-based spatial transcriptomics (ST) techniques have been groundbreaking in dissecting cell-cell communications within tissues by profiling positional gene expression. However, the most widely used ST technique, Visium Spatial Gene Expression by 10x Genomics (Visium), does not provide single-cell resolution, making it difficult to profile cell type-level information. Many reference-based deconvolution methods have been developed to increase its resolution, but the platform and batch effects between the reference and ST data compromise their accuracy. Here, we propose a new approach, **Re**gion-based cell **Sort**ing (ReSort), that generates a pseudo-internal-reference to reduce these platform effects. By simulating ST datasets under various scenarios, we demonstrate that ReSort significantly improves the accuracy of six state-of-the-art reference-based deconvolution methods. Moreover, applying ReSort to a mouse breast cancer tumor bearing both epithelial and mesenchymal clones identifies the spatial differences of immune cells between the clones, providing important insights for understanding the relationship between epithelial-mesenchymal transition and immune infiltration in breast cancer.

## BACKGROUND

Sequencing-based spatial transcriptomics (ST) techniques have been groundbreaking in dissecting cell-cell communications within tissues by profiling positional gene expression^1^. Researchers have applied this method to various diseases, including Alzheimer’s disease^2^, breast cancer^3^, prostate cancer^4^, melanoma^5^, and more. However, analyzing these datasets remains challenging due to the multi-cell resolution of the sequenced spatial units (referred to as spots). ST by Visium improved resolution to about five cells per spot, but each spot is still a mixture of different cells. To reveal the spatial organization of cell types and their changes with respect to other biological factors, such as pathology, it is critical to dissect cell type composition at every spot.

In addition to traditional bulk RNA-sequencing deconvolution techniques, such as CIBERSORTx^6^ and MuSiC^*7*^, many algorithms have been created to deconvolute cell type compositions for ST data. Stereoscope^8^ measures the cell types’ proportions by modeling cell type-specific gene expression as a negative binomial. Its parameters are learned from the single-cell RNA-sequencing reference data with cell type annotations and then applied to infer cell types’ proportions at every spot using maximum likelihood estimation. SPOTlight^9^ learns topic profiles from the reference data through non-negative matrix factorization. It then uses the learned topic-specific gene expression to estimate every spot’s cell type proportion through non-negative least squares. Another method, SpatialDWLS,^10^ first statistically assesses the enrichment of cell types and then applies a dampened weighted least square method to estimate the proportions of statistically enriched cell types.

All the aforementioned methods rely heavily on reference data from either single-cell RNA sequencing or a signature gene expression matrix, from which they learn cell type-specific gene expression to estimate the ratios of different cell types at every location in ST data. However, deconvolution becomes challenging if a proper single-cell reference is unavailable or is missing cell types. Moreover, technical noise such as batch and platform effects will affect the accuracy of the deconvolution results, especially when the references are from other experiments.

To address these problems, some approaches model platform-specific parameters to adjust for them. For example, RCTD^11^ and cell2location^12^ estimate gene-based platform effects in Bayesian models. Other methods, such as BayesSpace^13^, first detects spot-level clusters and uses spot-level information to estimate sub-spot compositions. It circumvents the platform effects within external references by generating a pseudo-internal reference. However, some spot-level clusters from BayesSpace might contain spots with mixed cell types, introducing noises for subspot-level estimations.

To address this problem, we propose a **Re**gion-based cell type **Sort**ing strategy (ReSort) that creates a pseudo-internal reference by extracting primary molecular regions from the ST data and leaves out spots that are likely to be mixtures. By detecting these regions with diverse molecular profiles, we can approximate the pseudo-internal reference to accurately estimate the composition at each spot, bypassing an external reference that could introduce technical noise.

In this study, we simulated ST datasets under different scenarios to demonstrate the efficacy of ReSort, finding that it generally improved the reference-based deconvolution methods’ accuracy by reducing technical effects. We also applied ReSort to a breast cancer ST sample to demonstrate its utility in investigating tumor microenvironments and tumor-infiltrating immune cells.

## RESULTS

### ReSort pipeline provides batch effect-free references for ST deconvolution

Most state-of-the-art deconvolution methods require single-cell RNA-sequencing data as references, but these references may introduce technical noise due to batch and platform differences. For example, single-cell RNA-sequencing of two pancreatic ductal adenocarcinomas (PDAC) tumor samples showed significant heterogeneity between individuals (Fig. 1a–b).

**Figure 1.**
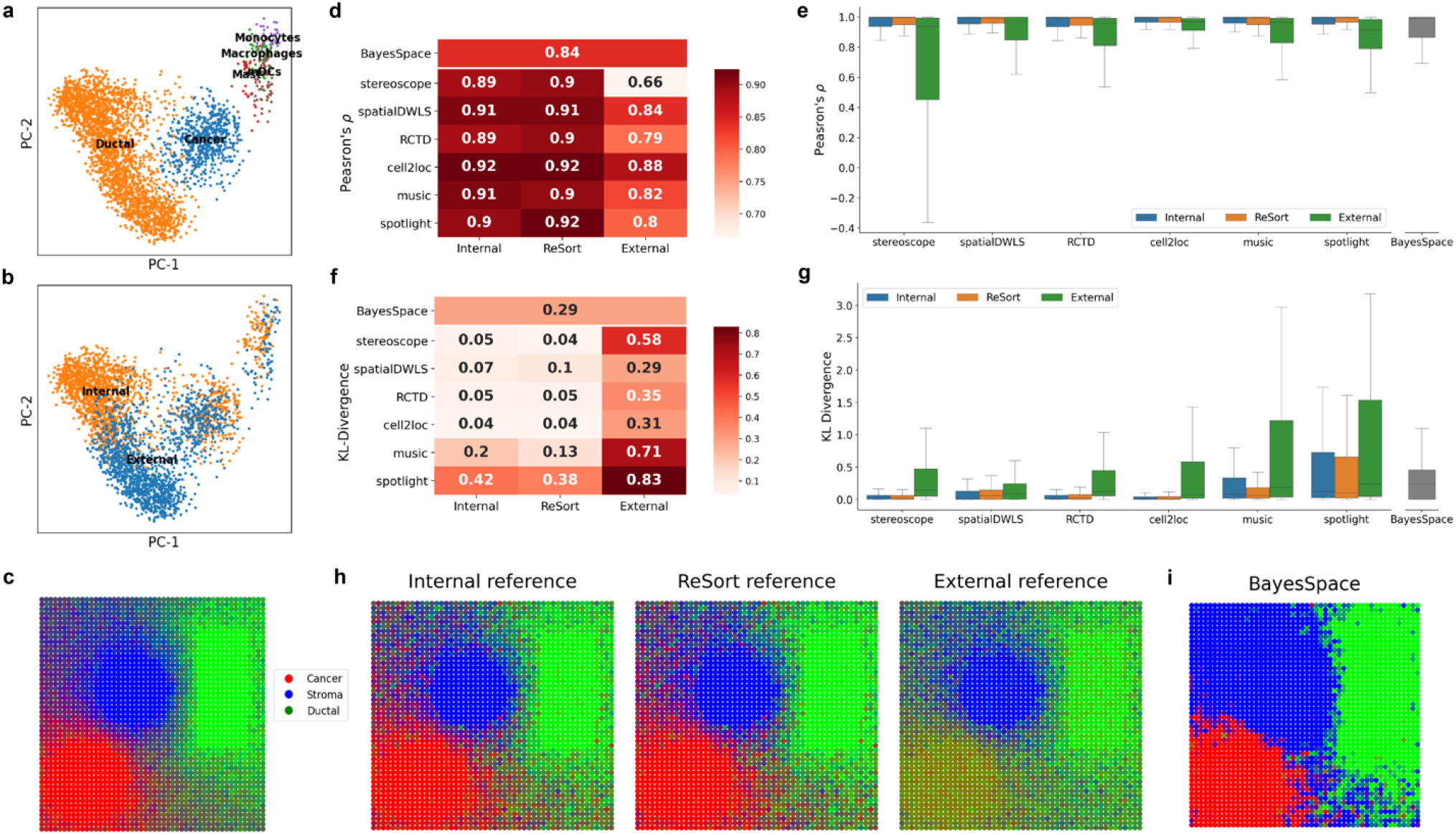
Benchmarking deconvolution methods’ performance on the ST sample simulated by primary cell types. **a-b** PCA visualization of two PDAC single-cell RNA-seq data colored by cell type (a) and reference type (b). **c** Visualization of simulated cell type proportions in RGB channels. **d-g** Performance comparison evaluated by Pearson’s correlation coefficient (d-e) and KL divergence (f-g). Heatmaps (d, f) show average metrics with rows as deconvolution methods and columns as the reference for each technique. Heatmaps’ increasing intensities of redness indicate higher raw values. Box plots (e, g) show the distribution of performances across the spots with columns as deconvolution methods. Box plots are defined with center line (median), box limits (upper and lower quartiles) and whiskers that extend at most 1.5 times of the interquartile range. **h** visualization of RCTD estimated cell-type proportions as an RGB image using different references. **i** Visualizing BayesSpace estimated cell-type proportions as an RGB image without reference. Spots colored in (c), (h) and (i) containing only one cell type will have pure red (cancer), blue (stroma), or green (ductal) colors, while spots with cell-type mixtures will have admixed colors.

Here, we introduce the **Re**gion-based cell type **Sort**ing (ReSort) strategy to address this issue. ReSort first extracts molecular regions from the ST sample as primary cell types using our published method, MIST^14^. Spots within each molecular region are highly similar to each other and likely to be dominated by one primary cell type, such as cancer. The advancement of ST by Visium provides high-resolution spatial profiles with thousands of spots, allowing ReSort to reliably leverage the detected molecular regions, each of which contains considerable number of spots, to form a pseudo-internal reference.

ReSort also allows the decomposition of finer cell types (referred to as secondary cell types), such as macrophages, that will not form a topological region in ST samples. Specifically, ReSort applies a two-step approach: it first deconvolutes the primary cell types, then estimates the proportion of secondary cell types through normalizing external-reference estimated secondary cell types’ proportions by the primary class proportion (see METHODS). ReSort thus constrains estimations of secondary cell types by their primary class proportion and reduces errors.

### ReSort accurately estimated primary cell type compositions

*In silico* simulation has been widely used to assess the efficacy of computational approaches by simulating faithful ground truth, a method we previously used to benchmark cell type deconvolution methods for bulk RNA-sequencing data^15^.

Here, we simulated ST data with 2,500 pseudo-spots on 50 by 50 grids (Fig. 1c, Supp. Fig 1a.) using a public pancreatic ductal adenocarcinomas (PDAC) single-cell RNA-sequencing data from one patient^16^. This data set was referred to as the internal reference since it is an ideal reference for deconvolution. However, such a reference does not exist in real deconvolution tasks. Thus, an external reference is often used by reference-based deconvolution methods. The single-cell RNA-sequencing data from another PDAC patient was introduced as an external reference to simulate actual deconvolution scenarios.

We first benchmarked the accuracy of six state-of-the-art deconvolution methods, including stereoscope^8^, spatialDWLS^10^, RCTD^17^, cell2location^12^, MuSiC^7^, and SPOTlight^9^, on three primary cell types—cancer, ductal, and stroma—with the external reference, internal reference, and the ReSort strategy. All reference-based approaches were further compared with a reference-free approach, BayesSpace.

To quantify the correlation between the estimated and ground truth proportions of cell types in each spot, we calculated Pearson’s correlation coefficient, referred to as ρ (Fig. 1d–e). We found that ReSort significantly increased ρ by 0.12, on average, compared to the external reference (p < 1e-10, two-sided paired t-test, N=2500). When compared to the internal reference, the average difference in ρ was negligible (Δρ = −0.004).

To assess the spot-level distributional divergence between the estimated and ground truth proportions of cell types, we calculated the relative entropy, also referred to as Kullback–Leibler (KL) divergence or D_KL_ (Fig.1 f–g). On average, ReSort achieved 80% lower D_KL_ than the external reference (p < 1e-10, two-sided paired t-test, N=2500) and 30% lower D_KL_ than the internal reference (p < 1e-10, two-sided paired t-test, N=2500). When compared to BayesSpace, ReSort achieved higher *ρ* in all cases and lower D_KL_ in five out of six cases.

To compare the estimated spatial patterns using different references, we transformed the cell types’ proportions into red-, green-, and blue-(RGB) colored channels where the strength of red, blue, and green indicates the abundance of tumor, stroma, and ductal cells, respectively (Fig. 1h, Supp. Fig. 1b). ReSort showed a similar RGB image to those of ground truth and the internal reference (Fig. 1a–b). In contrast, the external reference created an inaccurate pattern, especially in the tumor region. This result may have been caused by the heterogeneity in ductal and tumor profiles between the internal and external references (Fig. 1a–b), highlighting the challenges of using an external reference when deconvoluting ST data.

When comparing different reference-based approaches, we observed that although all methods achieved a similar level of Pearson’s correlation coefficients, stereoscope and RCTD had much lower divergence than other methods when using either an internal reference or ReSort. When using an external reference, spatialDWLS achieved the highest Pearson’s correlation coefficient (0.84) and slightest KL divergence (0.29) (Fig. 1). Notably, our findings were concordant with another benchmarking study^18^, which showed that cell2location, spatialDWLS, RCTD, and stereoscope significantly outperformed SPOTlight.

### ReSort identified immune infiltration in a simulation study

ST samples might pose greater challenges than the scenario described above. For example, tumor samples may be infiltrated by immune cells that are enclosed in the tumor. To simulate such cases, we randomly added immune cells to 10% of the tumor spots to mimic immune infiltration (Supp. Fig. 2). We then ran all the deconvolution methods previously mentioned with the three references: internal, external, and ReSort, respectively.

Compared with the external reference, ReSort consistently improved all the deconvolution methods’ Pearson’s correlation coefficients by 0.2 on average (p < 1e-10, two-sided paired t-test, N=2500) and decreased KL divergences by 87%, (p < 1e-10, two-sided paired t-test, N=2500) (Fig. 2a–b). Using the internal reference, ReSort significantly reduced the KL divergence by 32% (p < 1e-10, two-sided paired t-test, N=2500) while having similar average ρ values (Δρ = 2e-4, p=0.28, two-sided paired t-test, N=2500) (Fig. 2c–d). Importantly, BayesSpace failed to identify immune infiltration, with substantially lower ρ and higher KL divergence scores than any other approach (Fig. 2a–d). This may be because it overemphasized the spatial information in detecting spatial clusters (Supp. Fig. 2c), which will largely affect the successive and dependent sub-spot detection process.

**Figure 2.**
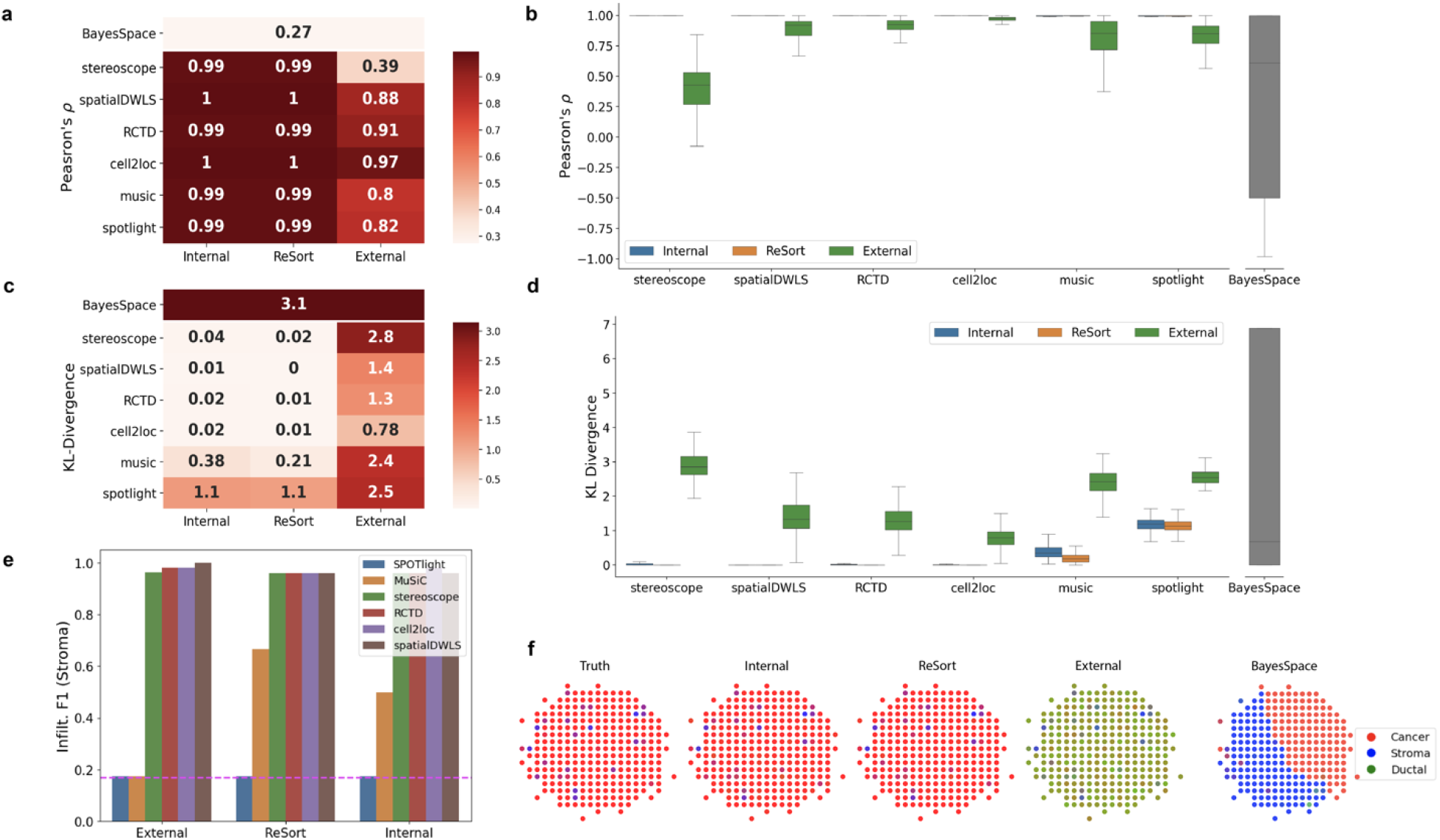
Benchmarking deconvolution methods’ performance on the ST sample with stroma infiltrated tumor spots simulated by primary cell types. **a-d** performance comparison evaluated by Pearson’s correlation coefficient (a-b) and KL divergence (c-d). Heatmaps (a, c) show average metrics with rows as deconvolution methods and columns as the reference for each technique. Heatmaps’ increasing intensities of redness indicate higher raw values. Box plots (b, d) show the distribution of performances across the spots with columns as deconvolution methods. Box plots are defined with center line (median), box limits (upper and lower quartiles), and whiskers that extend at most 1.5 times the interquartile range. **e** F-1 score for classifying stroma-infiltrated tumor spots. The X-axis represents the reference types used by deconvolution methods, pictured by bars with different colors. **f** From left to right: visualization as an RGB image of the cell type proportions in the cancer region using ground truth, RCTD with internal, ReSort, external references, and BayesSpace. Spots containing only one cell type have pure red (cancer), blue (stroma), or green (ductal) colors, while spots with cell type mixtures have admixed colors.

To assess various approaches’ abilities in detecting immune infiltrating events, we assessed their performances in classifying two classes of tumor spots: infiltrated tumor spots and pure tumor spots. Due to the imbalanced number of spots in each class, we evaluated precision, recall, and F1 score for each class. We also calculated a weighted F1 score to combine the F1 scores from both classes with weights defined by their relative sizes. Figure 2e shows that ReSort achieves an F1 score similar to the internal reference and outperforms the external reference. Interestingly, spatialDWLS, stereoscope, and RCTD showed much higher F1 scores than MuSiC and SPOTlight.

Additionally, the external reference failed all deconvolution methods in classifying pure tumor spots, resulting in lower F1 scores in the pure cancer class (Supp. Table 1., Fig. 2f). These results further highlight that batch effects impaired these methods’ ability to distinguish between ductal and cancer cell types (Fig. 1a–b). It also suggests ReSort’s utility in reducing the technical variation that may have existed in the external reference. In addition, MuSiC tends to predict more false-infiltrated spots, resulting in a lower recall score when classifying pure tumor spots and lower precision score when classifying immune-infiltrated spots. However, ReSort helped reduce MuSiC’s errors with a higher weighted F1 of 0.55 compared to 0.14 using the internal reference (Fig. 1a–d).

### ReSort accurately estimated secondary cell types’ compositions

Correctly analyzing the ST samples by their primary cell types provides important pathological information, but it is also critical to profile explicit cell types, such as immune subtypes in a pathological tissue sample, especially in cancer studies. To achieve this goal, ReSort applies a two-step strategy. ReSort first estimates the proportions of primary cell types, setting limits for the secondary cell types, such as macrophages, whose primary cell type is the stroma. Then, ReSort uses an external reference with only cell types belonging to each primary cell type—e.g., stroma—to estimate the relative abundance of individual secondary cell types.

The final estimation for the secondary cell types is calculated by the product of the proportion of primary cell types and the relative proportion of the secondary cell types (METHOD: eq. 5). Each primary cell type’s proportion serves as a precise prior for the secondary cell types, reducing potential errors in the estimated cell type proportions.

We used the same simulated immune-infiltrated PDAC sample to evaluate ReSort’s performance in the secondary deconvolution. Here, we excluded BayesSpace in benchmarking experiments due to its inability to estimate secondary cell types’ proportions.

When using ReSort as compared to the external reference, we observed an 18% increase in Pearson’s correlation coefficient (p < 1e-10, two-sided paired t-test, N=2500) and 66% decreased KL divergence (p < 1e-10, two-sided paired t-test, N=2500) (Fig. 3a–d). Moreover, the performance of all methods with an external reference resulted in much larger variations than using the internal reference or ReSort (Fig. 3b, d), suggesting unstable performances when using the external reference.

**Figure 3.**
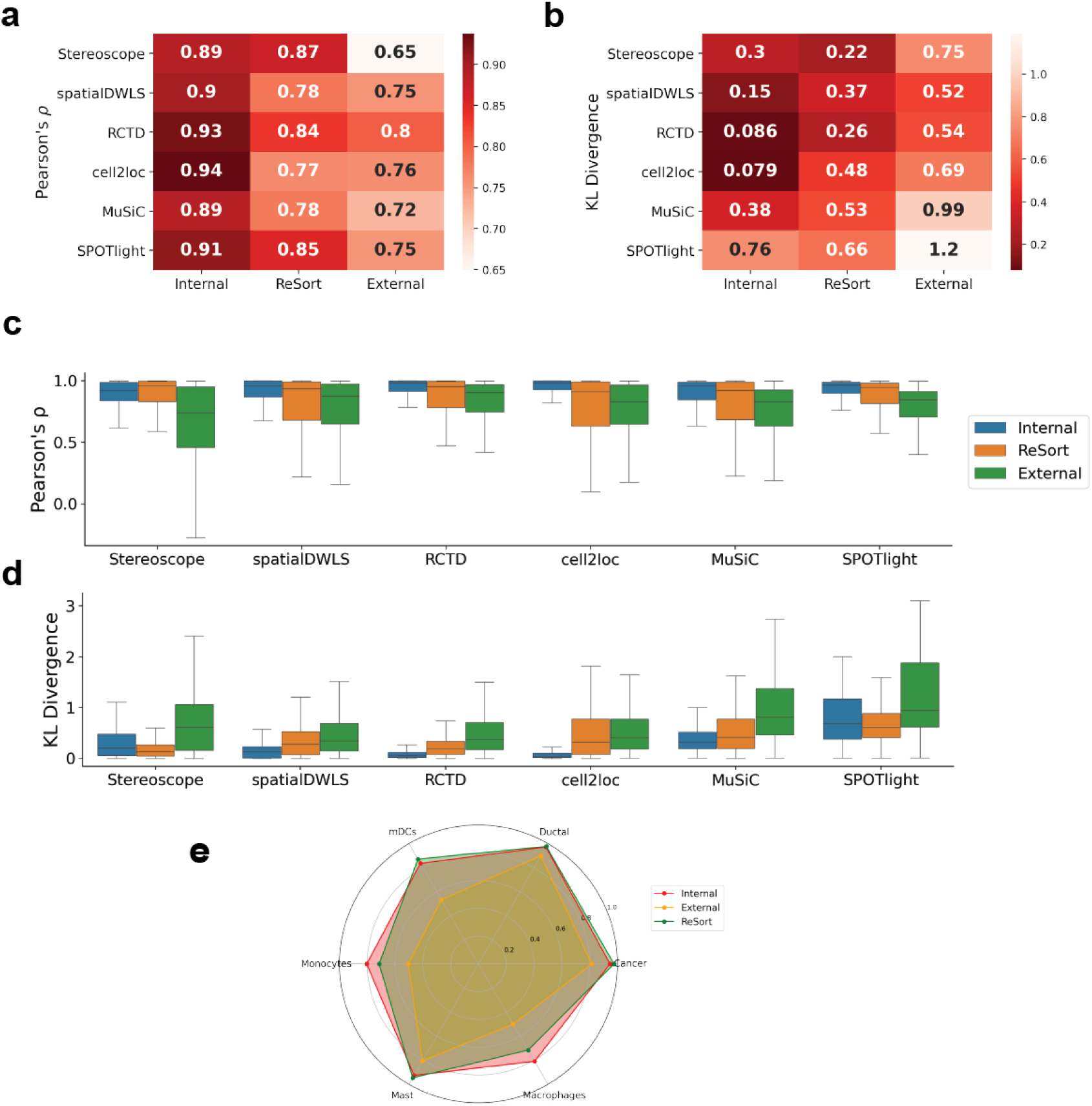
Benchmarking deconvolution methods’ performances on the ST sample simulated with rare cell types. **a-b** Heatmaps of deconvolution methods’ average performance in Pearson’s *ρ* (**a**) and KL divergence (b). **c-d** box plots showing distributions of deconvolution methods’ performance in Pearson’s *ρ* (c) and KL divergence (d). Box plots are defined with center line (median), box limits (upper and lower quartiles), and whiskers that extend at most 1.5 times the interquartile range. **e** Radar plot of Pearson’s *ρ* on each cell type using stereoscope with different references. *ρ* increases from 0 at the center to 1 on the outer loop. Each color represents a reference type. A larger area within the polygon indicates better overall performance.

With the external reference, RCTD had the highest Pearson’s correlation coefficient and the second smallest KL divergence when comparing with other deconvolution methods (Fig. 3). A potential explanation for this result is that RCTD was designed to reduce platform effect by estimating and removing gene-specific platform factors. When using ReSort, stereoscope achieves the highest Pearson’s correlation coefficient and the lowest KL divergence across all experiments (Fig. 3). This result suggests the importance of carefully selecting references for complex models like stereoscope, a deep learning-based model.

To further understand how ReSort improves stereoscope’s performance at the individual cell-type level, we calculated the Pearson correlation coefficient of each cell type between the estimated proportions and the ground truth proportions. We plotted performance as a radar plot with each direction representing a cell type, such that a better reference results in a larger area within the polygon (Fig. 3e). In doing so, we observed that the polygon generated using ReSort completely enclosed the polygon generated using the external reference, indicating that ReSort improves stereoscope’s performance in all cell types (Fig. 3e).

### ReSort discovered immune differences in the epithelial and mesenchymal clones of mouse breast cancer tumors

Epithelial-to-mesenchymal transition (EMT) is an important biological process during which cells lose polarity and show decreased adhesion and increased motility^19^. In cancer development, tumor cells at the primary site undergo EMT to gain higher migratory and invasiveness to metastasize to secondary organs, leading to a worse prognosis in clinical studies^20–22^. Studies have shown that various infiltrating immune cells shape the tumor ecosystem at different EMT states, thus affecting treatment outcomes^23–25^. How tumors with different EMT states co-evolve with the microenvironment yet retain a relatively distinct ecosystem is very important to cancer treatment but is less well understood.

To better understand the process of EMT in breast cancer, we applied ReSort to a mouse breast tumor sample containing both epithelial and mesenchymal tumor clones. ReSort first detected three primary regions in the sample using the MIST algorithm^14^ (see Method), including an epithelial tumor (E) region, a mesenchymal tumor (M) region, and a stroma (S) region (Fig. 4a, Supp. Fig. 4). Since each region contained spots with high intra-region transcriptional similarities, using these spots as a pseudo-internal reference allowed for accurate estimation of the proportions of primary cell types across the sample.

**Figure 4.**
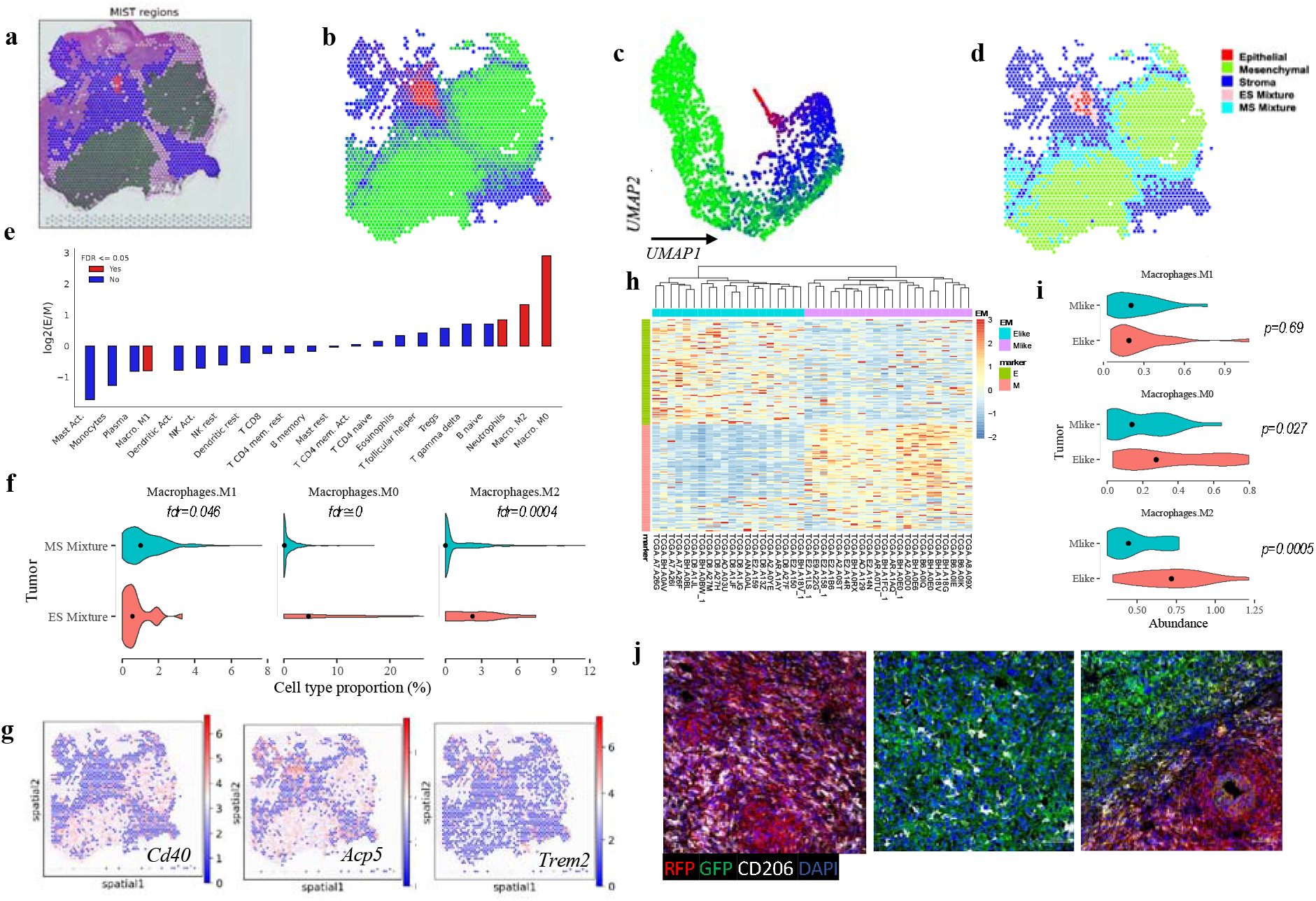
Applying ReSort to a breast cancer tumor with epithelial and mesenchymal tumor clones. **a** Molecular regions detected and mapped to the H&E staining image by MIST^14^. Red: epithelial clone; green: mesenchymal clone; blue: stroma clone. **b-c** Spatial (b) and UMAP (c) visualization colored by RGB channels using ReSort-estimated cell type proportions. **D** Stratifying spots into a pure epithelial clone (red), mesenchymal clone (green), stroma (blue), epithelial microenvironment (pink), and mesenchymal microenvironment (cyan). **e** Bar plot showing log2 of average fold changes (Y-axis, N=1 per bar) in LM22 (X-axis) immune cell types’ proportions in epithelial versus mesenchymal. **f** Violin plot comparing macrophage subtype’s proportions in epithelial and mesenchymal microenvironments (left to right: M1, M0, M2). Wilcoxon rank sum test used for statistical test with FDR adjusting for multiple comparisons. **g** *In situ* visualization of marker genes’ expression of macrophage subtype M1 (Cd40), M0 (Acp5), and M2 (Trem2). Expression values are colored from blue to red indicating increasing levels. **h** Epithelial-mesenchymal marker genes’ expression in 20 E-like and 22 M-like breast cancer tumors in The Cancer Genome Atlas (TCGA). **i** Violin plot comparing macrophage subtype’s proportions in E-like and M-like TCGA tumors (top to down: M1, M0, M2) with mean values marked as black dots. Statistical significances were derived using t-test without multiple-comparison adjustment. **j** Histochemical staining images. RFP: epithelial; GFP: mesenchymal; CD206: M2; DAPI: cell nucleus.

To validate the estimation’s accuracy, we colored the spots on the tissue and in the UMAP using the RGB channels representing the proportions of the corresponding primary cell types (Fig. 4b, c). Specifically, we used red for the epithelial tumor, blue for stroma, and green for the mesenchymal tumor. While pure E, M, and S spots formed clusters in both the H&E image and the UMAP, smooth transitions of colors were simultaneously observed at the boundary points in both figures (Fig. 4b, c). These results suggest that the ReSort-estimated primary cell types’ proportions represent both the physical and molecular structures of the data, preparing solid prior probabilities for secondary cell types’ deconvolution.

To estimate the roles of different immune cell types in the EMT process, we further stratified the admixture spots into epithelial-stroma (ES, N=32) and mesenchymal-stroma (MS, N=458) mixture spots (Fig. 4d), which represented the microenvironments of epithelial and mesenchymal tumors. Using a two-step ReSort strategy combined with CIBERSORTx^6^ and LM22 signature matrix, we estimated the proportions of 22 immune cell types in the ES and MS spots and statistically compared their compositions.

We observed that ES had 650% elevated M0 macrophages (p=8 x 10^-34^, t-test adjusted by multiple comparisons), 150% enriched M2 (p = 4e-4, t-test adjusted by multiple comparisons), and 179% increased neutrophils (p = 0.046, t-test adjusted by multiple comparisons). In contrast, MS had 174% elevated M1 macrophages (p=0.046, t-test adjusted by multiple comparisons) (Fig. 4e, f). Accordingly, we found significantly enriched expression of M0’s marker gene Acp5 (fold change = 150%, p = 7e-8, Wilcoxon rank-sum test) and M2’s marker gene Trem2 fold change = 68%, p = 7e-6, Wilcoxon rank-sum test) in the epithelial microenvironment (Fig. 4g). In contrast, Cd40, an M1 marker^26^, was significantly activated within the mesenchymal microenvironment (fold change = 146%, p = 5e-4, Wilcoxon rank-sum test) (Fig. 4g).

Using triple-negative breast cancer tumors (TNBC) from The Cancer Genome Atlas (TCGA), we validated that M0 and M2 were more enriched in epithelial-like TNBC tumors. We first extracted epithelial-like (N=22) and mesenchymal-like (N=20) TNBC tumors from TCGA^27^ using marker genes obtained from the ST sample (Fig. 4h). We then used CIBERSORTx^6^ to estimate the abundance of immune cell types in each tumor. We observed significantly enriched M0 (p=0.03, Wilcoxon rank-sum test) and M2 (p = 7e-4, Wilcoxon rank-sum test) in epithelial-like TNBC tumors (Fig. 4i). However, we did not observe significantly enriched M1 in the mesenchymal-like TNBC tumors (p=0.69, Wilcoxon rank-sum test).

To further validate our findings, we performed histochemical staining of mouse breast cancer tumors with both epithelial and mesenchymal clones. Specifically, we used CD206 to stain M2 cells and CD40 for M1 cells. Due to a lack of M0-specific cell surface marker, we didn’t stain M0. Both epithelial and mesenchymal clones showed M2 signals when cultured separately (Fig. 4j, left and middle), but we observed a substantially increased amount of M2 signals in the epithelial clone when both clones were cultured together (Fig. 4j, right). M1 staining was not enriched in the mesenchymal clone (Supp. Fig. 3).

### ReSort facilitates the discovery of tumor-infiltrating immune cells in mesenchymal mouse breast cancer tumors

Advancement of spatial transcriptomics techniques enables the profiling of spatial cell type enrichments, which is not possible in either bulk RNA-sequencing or single-cell RNA-sequencing datasets. Our ReSort pipeline, in contrast, was able to take advantage of this spatial information to research an important biomedical topic—i.e., tumor infiltrating immune cells, which play a critical role in disease outcomes^28,29^.

To understand tumor-infiltrating immune cells as compared with peripheral immune cells in the mesenchymal type of mouse breast cancer tumors, we further categorized MS spots as infiltrated or peripheral based on their location in the tissue sample, separated by the mesenchymal tumor region’s boundary as detected by MIST^14^. Infiltrated MS spots are found within the tumor while peripheral MS spots reside in the peripheral area outside the tumor (Fig. 5a).

**Figure 5.**
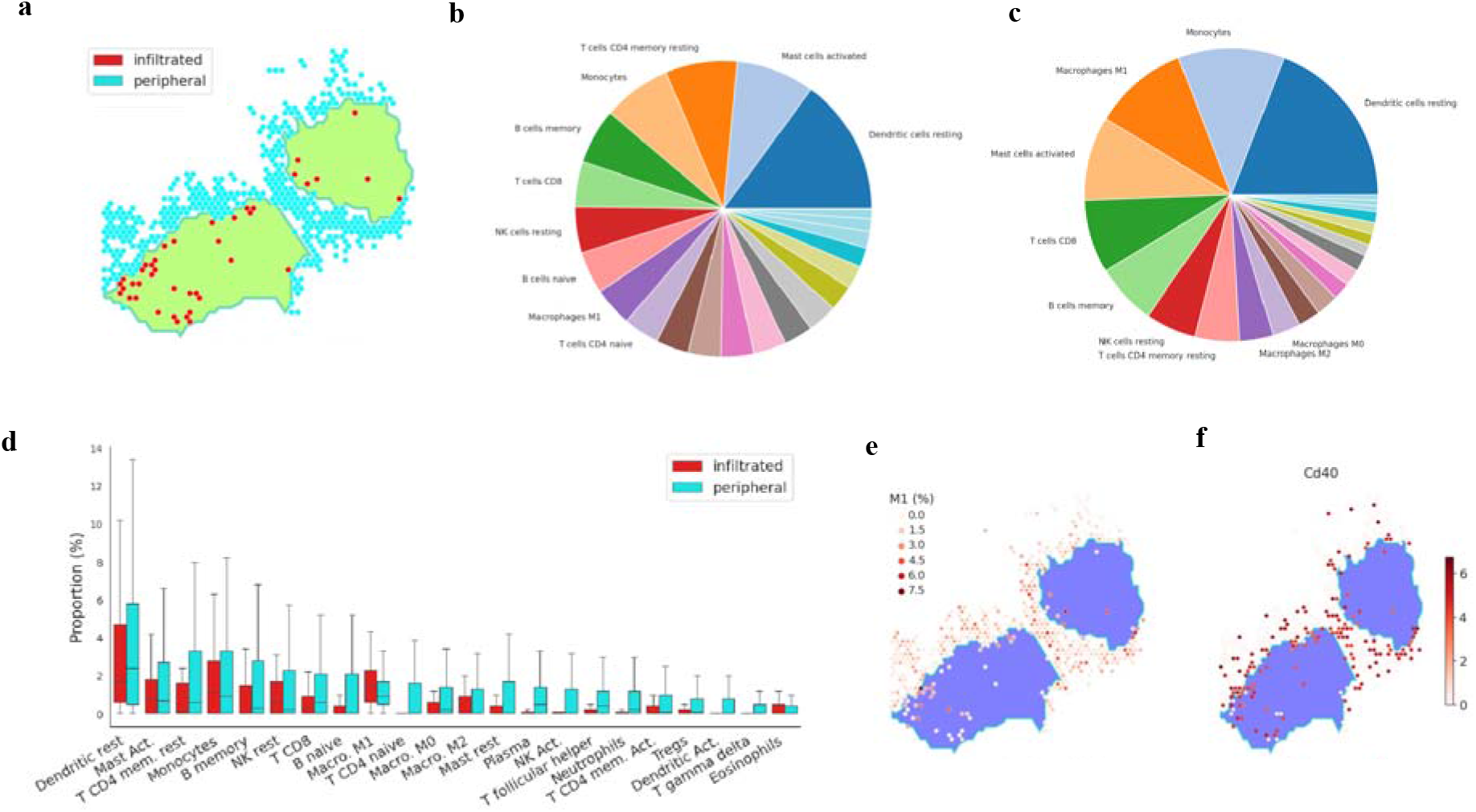
Applying ReSort to investigate immune infiltration in the mesenchymal clone of the breast cancer tumor. **a** Stratifying mesenchymal-stroma mixed spots into peripheral microenvironment (cyan) and immune-infiltrated tumor spots (red) separated by the boundary detected by MIST. **b-c** Pie charts of LM22 immune cell types’ relative proportions in the peripheral (b) and infiltrated (c) immune-tumor mixed spots estimated by ReSort. **d**: Box plots of LM22 (x axis) immune cell types’ proportions (y axis) estimated by ReSort. Box plots are defined with center line (median), box limits (upper and lower quartiles), and whiskers that extend at most 1.5 times the interquartile range. **e-f** *In situ* spatial visualization of M1 macrophage’s proportions (e) and its marker gene Cd40’s expression (f) in the mesenchymal microenvironment where the density of redness indicates higher values.

We then compared the most abundant immune cell types and found that naïve B-cells (rank=8) and naïve CD4 T cells (rank=10) ranked within the top 10 of the peripheral tumor microenvironments but were not in the top 10 of the infiltrated spots (Fig. 5b–c). Moreover, the relative abundance rank of M1 macrophages increased more in the infiltrated spots (rank=3) than in the peripheral spots (rank=9) (Fig. 5b–c).

Notably, M1 macrophage was the only immune cell type with significantly higher proportions in the infiltrated versus peripheral MS spots (fold change=42%, p = 0.017, t-test adjusted by multiple comparisons) (Fig. 5d, e). We observed that most infiltrated spots highly expressed M1’s marker gene, Cd40^26^, while many peripheral spots did not express it at all (Fig. 5f). This result suggests that M1 macrophage is more actively recruited into the tumor center rather than being gated by the boundary, like other immune cell types. Since M1s are anti-tumor macrophages^26^, we believe further investigation of their role in human EMT samples is critical to understanding and facilitating treatment.

## DISCUSSION

Precisely estimating the composition of cell types in spatial transcriptomics (ST) is critical to understanding their communication in the tissue. Currently, most state-of-the-art deconvolution approaches use an external reference to estimate the proportion of cell at every spot. However, using an external reference may cause technical noise due to batch and platform effects, leading to inaccurate results. In this study, we designed a **Re**gion-based digital cell **Sort**ing (ReSort) strategy for sequencing-based spatial transcriptomics (ST) data sets. ReSort learns a pseudo-internal reference from the ST spots by detecting core regions, each containing a group of spots with high transcriptome-level similarities. It can then use reference-based deconvolution methods to increase their accuracy by reducing batch and platform effects.

We first demonstrated that ReSort significantly improves other reference-based deconvolution methods’ accuracy in Pearson’s correlation coefficient score and KL divergence. To comprehensively evaluate our strategy, we simulated three scenarios: a base case with three primary cell types, a tumor sample with immune infiltration, and a finer (secondary) cell type estimation. In all three simulation studies, ReSort achieved significantly higher correlation and smaller KL divergence. Moreover, ReSort’s performance was not statistically different from the ideal-yet-nonexistent internal reference used to generate the simulation data. These simulation results thus demonstrate that ReSort can provide a pseudo-internal reference to increase reference-based deconvolution methods’ accuracy by avoiding batch and platform effects.

To understand immune cells’ roles in tumor-related epithelial-to-mesenchymal transition (EMT), we also applied ReSort to a mouse breast cancer sample with both epithelial and mesenchymal tumor subtypes. Surprisingly, we found polarization of macrophages in different tumor clones. Specifically, we found M1 macrophage significantly enriched in the mesenchymal tumor clone’s microenvironment, while M2 and M0 were substantially elevated in the epithelial tumor clone’s microenvironment. Additional validation in TCGA triple-negative breast cancer tumors and histological staining of CD206 confirmed M2 and M0’s enrichments in the epithelial-like tumors. Moreover, by separating the tumor-immune mixtures in the mesenchymal clone, we found that M1 was the only tumor-infiltrating immune cell type with a higher average proportion in the tumor center than the peripheral microenvironment.

Although ReSort’s findings on the mouse breast cancer sample revealed novel insights for understanding immune cells’ functions in EMT, the lack of validation on human ST samples is a limitation of our study. Since many other studies have reported that tumor-infiltrating immune cells and tumor-associated macrophages are critical to the outcome of immune therapies^28,30^, we believe further validation using human samples is essential to provide greater insights for treatment.

## METHODS

### Single-cell RNA-sequencing data collection

Single-cell RNA-sequencing of pancreatic ductal adenocarcinomas was obtained from Gene Expression Omnibus under accession number GSE111672. We used the filtered raw count matrix of cells from PDAC-A as building blocks to simulate RNA profiles of spatial transcriptomes and referred to it as the internal reference. In contrast, the single-cell RNA-sequencing data from the other patient (PDACA-B) used in the same study^16^ was referred to as the external reference.

The PDAC-A and the PDAC-B samples shared six cell types: cancer, ductal, macrophages, mast, mDCs, and monocytes. We further categorized them into three primary cell types, with macrophages, mast, mDCs, and monocytes grouped as stroma.

### Simulating patterns of cell types

We first generated a tissue grid of 2,500 spots with a layout of 50 rows and 50 columns. To mimic the pathological organization of tissues, three major regions with homogeneous cell populations—namely, the cancer, ductal, and stroma regions—were simulated. The cancer and stroma regions were circular and the ductal region was rectangular (Fig. 1).

To simulate the ratio of cell type **r** at every out-of-region spot *s*, we denoted the affinity A_*S*,**r**_ between spot *s* and region **r** as the inverse of the Euclidean distance between *s* and the closest point, *s*_**r**_ within each region **r**, with a random Gaussian noise, *δ*_**r**_ (Equation 1). We then normalized the proportions to sum to one (Equation 2).

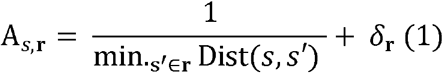

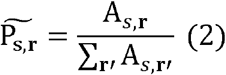

Since every spot in Visium ST data captures about ten cells, to transfer cell type proportions to counts we assigned the count of each cell type **r**, denoted by C_s,**r**_, by multiplying the ratios obtained in Equation (2) by ten and taking the integer values (Equation 3). The proportion values are then re-normalized and saved for later evaluation (Equation 4).

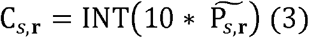

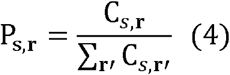

### Sample cells and reads to form count matrix

We sampled C_*s*,**r**_ cells from type **r** without replacement for every spot *s*. To make the library size of each spot in the simulated ST data comparable to that of actual Visium data sets, we down-sampled 10% of the gene counts from every sampled cell to keep the number of reads in every simulated spot at the same magnitude as the Visium data. All cells’ sampled gene counts were then aggregated to make a simulated mini-bulk gene expression count matrix.

### Simulate spots with cell infiltration

To simulate cases in which one cell type infiltrated a core region of other cell types—e.g., immune infiltration into the cancer area—we randomly sampled 10% of the spots in the cancer area and added stroma cells artificially. The proportion of stroma cells for every such spot was sampled from a Gaussian distribution (*μ* = 0.5, *σ* = 0.2). The resampled count matrix was then generated using the aforementioned steps (Equations 1 to 4).

### Extract primary molecular regions using MIST

We used a previously developed method, MIST^14^, to extract homogeneous regions to detect primary molecular regions. MIST combines the molecular similarity and the physical adjacency among spots to determine highly homogeneous regions in the sequenced tissue^14^.

MIST normalizes the count matrix by the library size and transforms the resulting gene expression values at a log scale. It then performs principal component analysis (PCA) on the normalized gene expression values and calculates a similarity score for spatially adjacent spots using the first *k* principal components. The similarity score serves as the weight of the edge *E* =< *u, v*> for spot *u* and *v*. MIST, then filters out edges with low weights using an automatically optimized threshold epsilon. Each connected component in the remaining graph, with more than *m* nodes, serve as the molecular regions, where *m* is a constant with the default value of 40 for Visium data.

To validate the accuracy of the annotations on our simulated sample, we calculated a Rand Index of 0.74 between annotated regions and the ground truth using the Python scikit-learn package. Supp. Fig. 1b shows the visual concordance between MIST-detected regions and the ground truth.

### Primary deconvolution

We define primary deconvolution as the task of estimating the proportion of primary cell types (regions) in the ST samples. In the simulation study, we grouped the six cell types in the PDAC-A and PDAC-B single-cell RNA-sequencing data into three primary cell types: cancer, ductal, and stroma.

For ReSort, this task does not require additional single-cell RNA-sequencing references and avoids technical effects when estimation is based on external references. A curated reference matrix **Ω** was generated. Every row in **Ω** is a spot with an annotated primary cell type, and every column is a gene. **Ω** contains sufficient source information required by all the compared deconvolution methods. For every spot, we denote 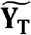 as the estimated proportion for primary cell type **T**, with all proportions summed to be one.

### Secondary deconvolution

We define secondary deconvolution as the task of estimating the proportions of finer cell types, such as macrophages. These finer cell types generally do not form topological structures in the ST samples, so they could not be directly extracted using MIST.

To achieve this goal, ReSort used a two-step deconvolution strategy. It first applied the primary deconvolution mentioned above to estimate the proportions of every primary cell type **T**’s proportion, denoted as 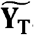. It then used an external reference ω—either single-cell RNA-sequencing data or a signature matrix, such as immune signature matrix LM22^6,31^—to estimate the secondary cell types’ proportions, 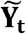, for each sub-type **t**. The final estimation of the proportion of subtype **t** is then calculated in Equation 5.

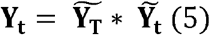

### Running deconvolution methods

We ran each deconvolution method listed below in two modes: primary and secondary. In the primary deconvolution mode, we used three references: internal, external, and ReSort. To ensure consistency and enable fair benchmarking, we used the same parameters for each tool when using different references.

The internal reference refers to the single-cell RNA-sequencing data (PDAC-A) used to generate the simulated ST data, which does not exist in reality. The external reference refers to the single-cell RNA-sequencing data from the same tissue but different samples (PDAC-B). The external reference represents the scenarios for most deconvolution tasks using the state-of-the-art methods listed below. The MIST reference is extracted from the ST data itself to avoid the technical effects caused by an external reference. The returned proportions of all cell types were further normalized to guarantee they were summed to one. All code needed to reproduce our results is available at GitHub (https://github.com/LiuzLab/ReSort_manuscript).

### Running RCTD

We ran RCTD using the commands instructed by the authors at https://github.com/dmcable/spacexr. Specifically, we used the function creat.RCTD() with ten cores, and the function run.RCTD() with parameter “doublet mode = full” to enable estimating for more than two cell types.

### Running MuSiC

We ran MuSiC by following the instructions (https://xuranw.github.io/MuSiC/articles/MuSiC.html) provided by the authors of the package. We used the music_prop() function with default parameters.

### Running stereoscope

We followed the authors’ instructions at https://github.com/almaan/stereoscope. We used default values provided by the instructions except for reducing the epoch values to reduce the computational time and avoid overfitting. Instead, we used an epoch value of 10,000 for each experiment in training the reference and the ST data, while the default value is 75,000.

### Running SPOTlight

We followed instructions to run the function spotlight_deconvolution(). Marker genes for every cell type were detected using the function FindAllMarkers() from the R package Seurat^32^. We normalized the raw count data by the library size and then transformed it into the log scale. Markers for each cell type were then extracted for genes with over 20% fold change and adjusted p-value of less than 0.05.

### Running SpatialDWLS

We used the function runSpatialDeconv() from the R package Giotto. Using the same procedure to extract marker genes when running SPOTlight, we generated a signature matrix by first detecting marker genes for each cell type. We used default values for all other parameters.

### Running cell2location

We followed the tutorial from https://cell2location.readthedocs.io/ to perform cell2location with our data. With the loading and basic preprocessing of the Visium and reference data, we fitted and extracted the cell type-specific expression signatures from the reference data using the RegressionModel() function with 800 epochs to ensure convergence. We then used the cell2location() function to estimate the abundance of cell types at each spot after training the cell2location model with 30,000 epochs. All the training was done in one GPU node and took 40-50 minutes.

### Running BayesSpace

We ran BayesSpace by following its tutorial at https://edward130603.github.io/BayesSpace/articles/BayesSpace.html.We constructed a SingleCellExperiment object using raw counts and spatial coordinates. We used default parameters with log-normalized gene expression data, the top 2,000 highly variable genes, and seven principal components. Next, we extracted spot-level clusters using the function spatialCluster() with 10,000 iterations, and subspot-level clusters using function spatialEnhance() with 100,000 iterations.

### Running CIBERSORTx

We ran CIBERSORTx^6^ with the library-size normalized count matrix without log-scale transformation, as suggested by Jing et al^15^. To run CIBERSORTx, we used the default LM22 reference data, with 1000 permutations.

### Evaluation of simulation methods

We used Pearson’s correlation coefficient, referred to as ρ, to estimate the concordance between the estimated cell types’ proportions and the ground truth (Equation 6).

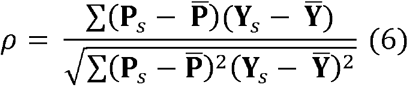

**P**_*s*_ denotes the ground truth spot *s*’s proportion and **Y**_s_ represents the estimated spot *s*’s proportion. The metric *ρ* is used to assess the estimated values’ accuracy at the given spot *s*. We used the function stats.pearsonr() from the Python package scipy to calculate ρ.

The second metric we evaluated is the Kullback–Leibler (KL) divergence, also referred to as relative entropy and D_KL_ (Equation 7).

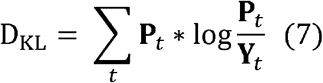

D_KL_ captures the divergence between the ground truth distribution and the estimated distribution of cell type compositions.

A good method should have a high ρ score and a low D_KL_ score.

Moreover, to assess the models’ performance in detecting immune infiltration in the tumor region, we calculated the precision, recall, and F1 scores for two classes: infiltrated and pure tumor spots. An infiltrated tumor spot is a spot in the tumor region with a stroma proportion of no less than 5%. A pure tumor spot is a spot in the tumor region with a tumor proportion greater than 95%. With the definition of these two classes, we calculated the precision, recall, and F1 scores for each class (Equation 8–10).

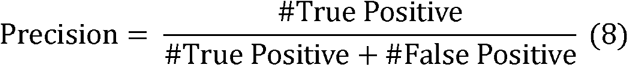

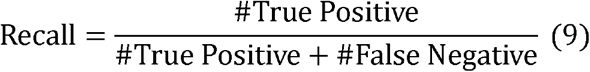

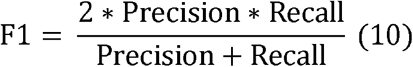

We systematically evaluated the classification accuracy by showing each class’s precision, recall, and F1 scores (Supp. Table 1). Additionally, we calculated a weighted F1 score defined as F1_weighted_ = *α* * F1_pure_ + (1 - *α*) * F1_infilt_, where 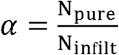, with N_pure_ representing the number of pure cancer spots and N_infilt_ defining the number of immune-filtrated spots within the tumor region.

### EMT Spatial Transcriptomics experiments

PyMT-N and PyMT-M cells were derived from the spontaneous tumor of the MMTV-PyMT mouse model to represent epithelial and mesenchymal types, respectively. PyMT-N and PyMT-M cells mixed at a 1:1 ratio (2.5×105 cells/each) were resuspended in Matrigel and injected into the #4 mammary fat pads of C57/B6 mice. On day 21 after injection, the tumors were resected at approximately 1 cm after the mice were euthanized and perfused with PBS. The tumors were then excised and embedded with OCT in an isopentane bath kept in dry ice. The blocks were then sectioned at 10 μm thick and placed on the designated regions of a 10x Genomics Visium Spatial Gene Expression slide. After H&E staining, each section was imaged using a bright color field by Cytation 5. The sections were then processed following 10X Genomics gene expression protocols until the libraries were constructed, which were sequenced by Novaseq 6000 with 150bp paired-end reads. About 300M reads were recorded for each sample.

We used the software SpaceRanger-1.3.0 to process the mouse breast cancer ST samples with both epithelial-type and mesenchymal type tumors on the samples. After running SpaceRanger, the filtered count matrix was used for further analysis.

### *Molecular region detection and annotation for the PyMT-M-N* mouse breast cancer sample

We preprocess the sample with MIST^14^ using the function preprocess(species=“Mouse”, hvg_prop=0.9, n_pcs=10), which removed uninformative genes with low coverage and variance. We then called the function extract_regions(sigma=0.4, min_region=3, gap=0.02), which extracted at least 40% of all spots and assigned them to at least three major regions.

We molecularly annotate the PyMT-M-N mouse breast cancer sample that have both epithelial and mesenchymal clones using previously published epithelial- and mesenchymal-associated markers^33^. We used Krt18 and Fn1 as epithelial transcriptomics markers, and Zeb1 and Twist1 as mesenchymal markers to label the epithelial region and mesenchymal region. Supp. Fig. 3 shows that substantially higher read counts from the green and red regions indicate these two are likely to be tumor regions, where cells having higher densities and are more proliferated. Moreover, we observed higher expression values of Krt18 and Fn1 in the red region, while Zeb1 and Twist1 are enriched in the green region, suggesting that red region being epithelial tumor and green region as mesenchymal tumor. We assigned blue region as a stroma region because it has a low total number of gene counts and is not enriched with either epithelial or mesenchymal markers.

### External validation using epithelial and mesenchymal-like TCGA cancer tumors

We used the function getTCGA(disease=“BRCA”, data.type=“RNASeq”, type=“RPKM”) from the TCGA2STAT R package to obtain breast cancer RNAseq data^34^. We then extracted triplenegative breast cancer (TNBC) tumors using the meta information^27^, resulting in 128 tumor samples. We normalized the RPKM by the library size and performed log-transformation.

Then, we extracted top 100 epithelial-enriched marker genes and the top 100 mesenchymal-enriched marker genes from the ST samples to perform clustering on the 128 TNBC samples. Based on the clustering results from the R package pheatmap (Supp. Fig. 4), we extracted 20 epithelial-like and 22 mesenchymal-like TNBC tumors. We ran CIBERSORTx in the absolute mode to decompose immune cell types’ abundances in these 44 TNBC E- and M-like tumors.

### Immunohistochemistry staining on frozen primary tumors

Tumor-bearing mice were euthanized and perfused with 30mL PBS before the primary tumors were removed and embedded immediately in OCT in an isopentane bath pre-chilled to −80 □ and maintained on dry ice. Tumor sections at 10μm thickness were fixed with 10% neutral buffered formalin in PBS and permeabilized with 0.2% Triton X-100 in PBS for 10min at room temperature and blocked with 10% normal donkey serum in PBS-GT (PBS with 0.1% Triton X100 and 0.1% gelatin) for 1 hour at room temperature. Sections were then incubated with primary antibodies (chicken anti-GFP, 1:200, Novus Biologicals, NB100-1614; rabbit anti-RFP, 1:200, Rockland, 600-401-379; goat anti-CD206, 1:100, R&D Systems, AF2535) overnight in a humidified chamber at 4□. Slides were then washed and incubated with Alexa Fluor 488-conjugated donkey anti-chicken (1:400, Jackson ImmunoResearch), Alexa Fluor 555-conjugated donkey anti-rabbit (1:400, Jackson ImmunoResearch) and Alexa Fluor 647-conjugated donkey anti-goat (1:200, Jackson ImmunoResearch) secondary antibodies for 2 hours at room temperature in a humidified chamber. Sections were then washed and stained with Hoechst 33342 (Thermo Fisher Scientific 62249) and mounted with Prolonged Gold Antifade Mountant (Molecular Probe). Images were acquired by a Zeiss LSM780 confocal microscope.

## DATA AVAILALIBITY

Single-cell RNA-sequencing of pancreatic ductal adenocarcinomas was obtained from Gene Expression Omnibus under accession number GSE111672. Simulated ST and the mouse PyMT-M-N Visium ST sample were deposited at Zenodo (doi: 10.5281/zenodo.7434870)^35^.

## CODE AVAILABILITY

The code to reproduce the results in this manuscript is at: https://github.com/LiuzLab/ReSort_manuscript.

## ACKNOWLEDGEMENTS

Research reported in this publication was partially supported by the Eunice Kennedy Shriver National Institute of Child Health & Human Development of the National Institutes of Health under Award Number P50HD103555 for use of the Bioinformatics Core facilities. The content is solely the responsibility of the authors and does not necessarily represent the official views of the National Institutes of Health. Z.L., L.WANG, C.L., and W.W. are also partially supported by the Chao Endowment.

## AUTHOR CONTRIBUTIONS

L. WANG and Z. L. conceived the project. L. WANG developed and implemented the methods, collected the data, and designed and performed the computational experiments and analyses. W. W and C. L. involved partially in computational experiments for benchmarking. L. WU and X. Z. designed and performed biological experiments to validate the computational pipeline. Z. L. supervised the project and led the discussions about the results. L. WANG drafted the manuscript. All authors contributed to the final manuscript.

