## Supplementary Figures & Table 1 for "Accurate cell type deconvolution in spatial transcriptomics using a batch effect-free strategy"

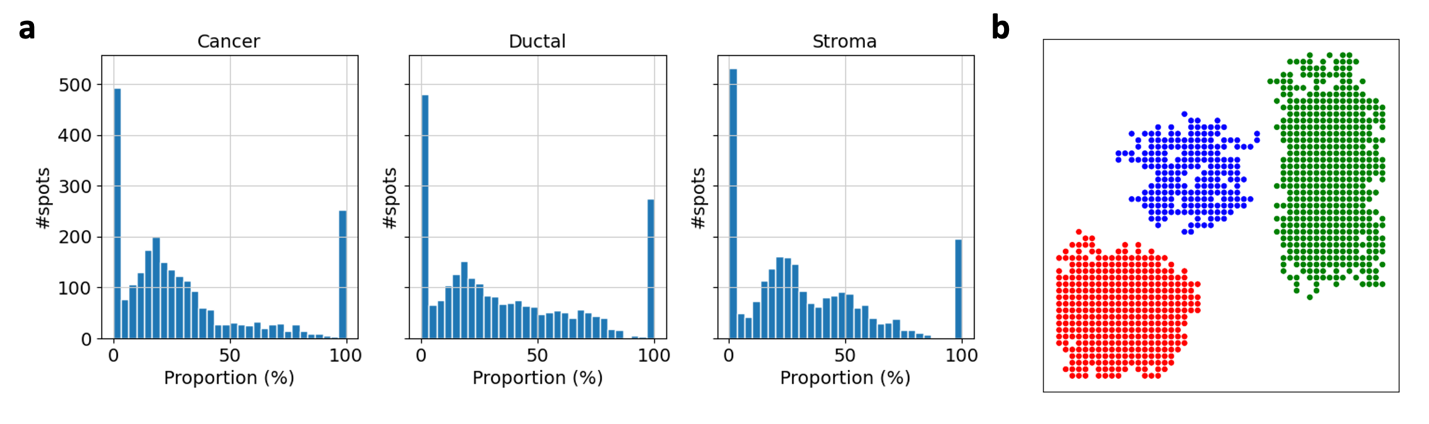

**Supplementary Figure 1** | Simulation of ST samples using primary cell types. **a**: Simulated proportional distributions of three primary cell types. **b**: MIST detected regional spots for three major cell types. Red: cancer; blue: stroma; green: ductal.

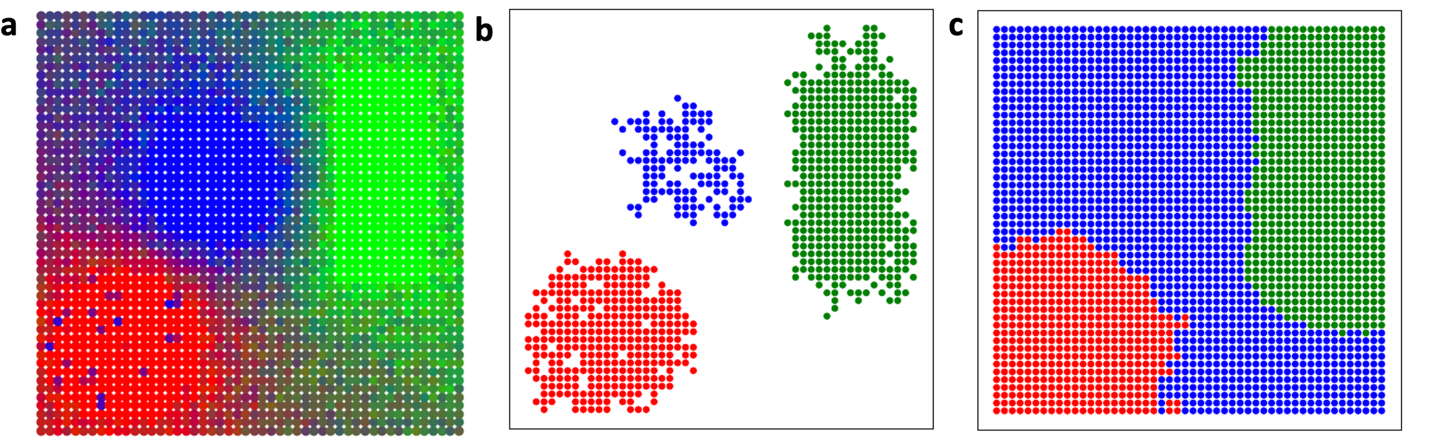

**Supplementary Figure 2** | Simulation of ST samples with stroma infiltrated tumor spots using primary cell types. **a**: Spatial pattern of the simulated ST sample. Each spot is colored by primary cell types’ proportions denoted by red as cancer, green as ductal and blue as stroma cell types. **b**: MIST detected regional spots for three major cell types. Colors are matched with (a). **c**: BayesSpace’s spatial clusters.

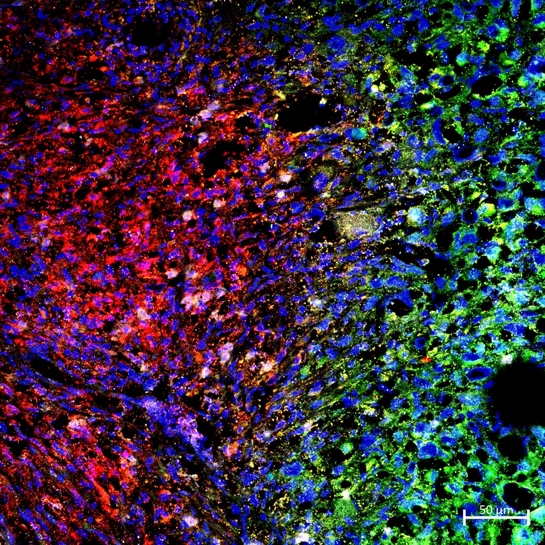

GFP RFP CD40 DAPI

**Supplementary Figure 3**| Histological staining of M1 using CD40 in breast cancer tumor with epithelial and mesenchymal clones. RFP: Epithelial; GFP: Mesenchymal; CD40: M1; DAPI: cell nucleus.

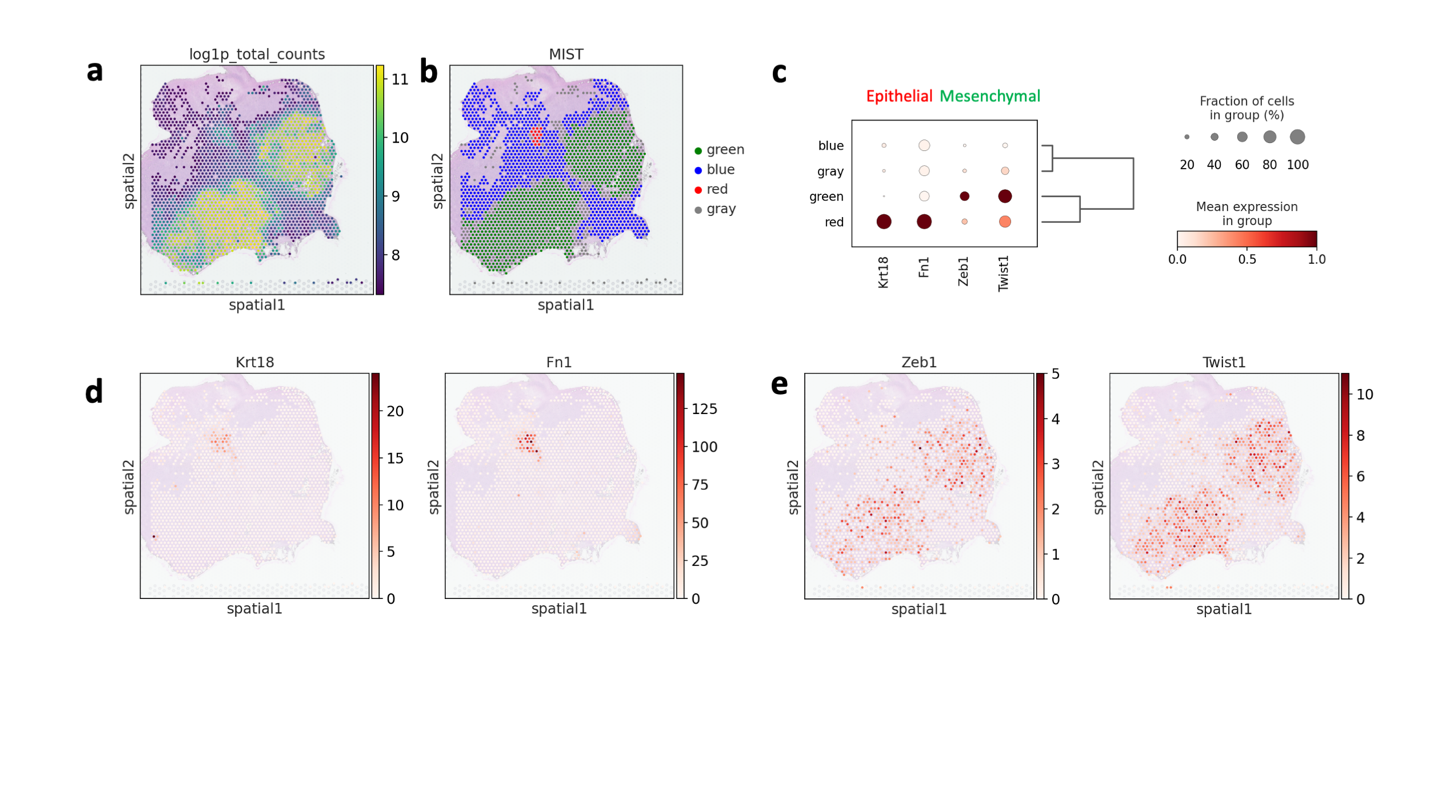

**Supplementary Figure 4** | **Annotating PyMT-M-N breast cancer tumor. a** Heatmap of log-scaled total number of gene counts in the sample. **b** regions detected by MIST in reen, blue and red colors. Gray colored spots are not assigned to either major regions. **c** Expression dot plot of epithelial (Krt18 and Fn1) and mesenchymal markers (Zeb1 and Twsit1). **d** Heatmap of epithelial marker genes’ (Krt18 and Fn1 expression with higher values colored in red. **e** Heatmap of mesenchymal marker genes’ (Krt18 and Fn1 expression with higher values colored in red.

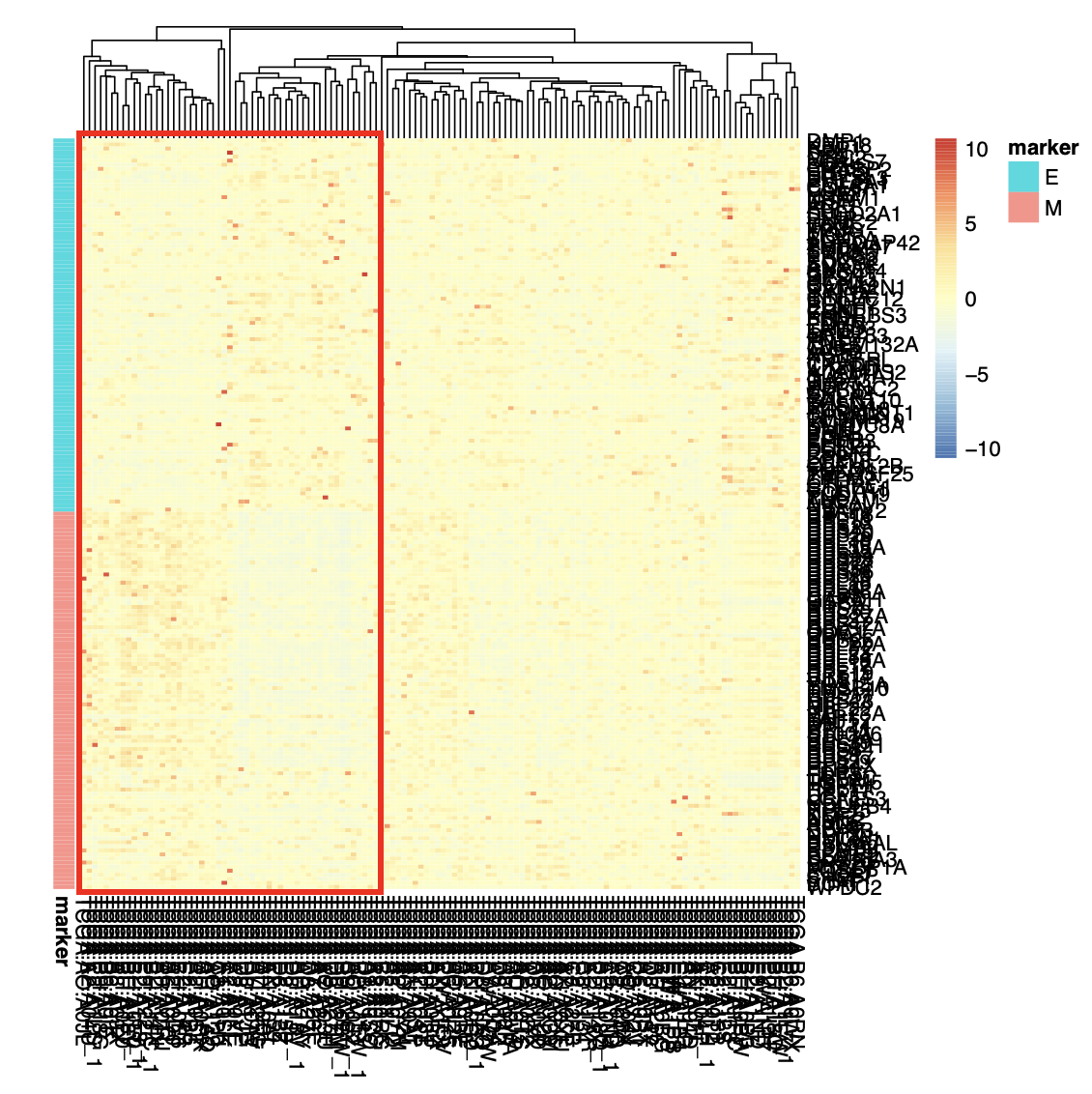

**Supplementary Figure 5** | Heatmap of epithelial and mesenchymal marker genes’ expression in 128 triple negative breast cancer (TNBC) tumors from TCGA. Genes are annotated by marker groups in the row annotations. Columns of the red box are the 44 epithelial- and mesenchymal-like TNBC tumors used for further study.

**Supplementary Table 1.** Classification report on pure tumor and tumor infiltrated stroma spots

| **Precision Cancer** | **Recall Cancer** | **F1 Cancer** | **Precision Stroma** | **Recall Stroma** | **F1 Stroma** | **F1 Weighted** | **reference** | **method** |
| --- | --- | --- | --- | --- | --- | --- | --- | --- |
| **0** | 0 | 0 | 0.095588235 | 1 | 0.174496644 | 0.016679826 | External | spotlight |
| **0** | 0 | 0 | 0.095588235 | 1 | 0.174496644 | 0.016679826 | Internal | spotlight |
| **0** | 0 | 0 | 0.095588235 | 1 | 0.174496644 | 0.016679826 | ReSort | spotlight |
| **0** | 0 | 0 | 0.095940959 | 1 | 0.175084175 | 0.016735987 | External | music |
| **0** | 0 | 0 | 0.961538462 | 0.961538462 | 0.961538462 | 0.091911765 | External | stereoscope |
| **0** | 0 | 0 | 1 | 0.961538462 | 0.980392157 | 0.093713956 | External | RCTD |
| **0** | 0 | 0 | 1 | 0.961538462 | 0.980392157 | 0.093713956 | External | cell2loc |
| **1** | 0.052845528 | 0.1003861 | 0.337837838 | 0.961538462 | 0.5 | 0.138584488 | Internal | music |
| **1** | 0.028455285 | 0.055335968 | 1 | 1 | 1 | 0.145634736 | External | spatialDWLS |
| **1** | 0.369918699 | 0.540059347 | 0.510204082 | 0.961538462 | 0.666666667 | 0.552161517 | ReSort | music |
| **0.908396947** | 0.483739837 | 0.631299735 | 0.101449275 | 0.538461538 | 0.170731707 | 0.58727485 | BayesSpace | BayesSpace |
| **0.991266376** | 0.922764228 | 0.955789474 | 1 | 0.923076923 | 0.96 | 0.95619195 | Internal | stereoscope |
| **0.995744681** | 0.951219512 | 0.972972973 | 1 | 0.961538462 | 0.980392157 | 0.97368216 | Internal | cell2loc |
| **0.991666667** | 0.967479675 | 0.979423868 | 1 | 0.923076923 | 0.96 | 0.977567175 | Internal | RCTD |
| **0.991735537** | 0.975609756 | 0.983606557 | 1 | 0.923076923 | 0.96 | 0.981350048 | ReSort | stereoscope |
| **0.991803279** | 0.983739837 | 0.987755102 | 1 | 0.923076923 | 0.96 | 0.985102041 | ReSort | cell2loc |
| **0.991869919** | 0.991869919 | 0.991869919 | 1 | 0.923076923 | 0.96 | 0.988823529 | Internal | spatialDWLS |
| **0.991902834** | 0.995934959 | 0.993914807 | 1 | 0.923076923 | 0.96 | 0.990672951 | ReSort | RCTD |
| **0.991935484** | 1 | 0.995951417 | 1 | 0.923076923 | 0.96 | 0.992514884 | ReSort | spatialDWLS |
